# Clonal analysis of fetal hematopoietic stem/progenitor cell subsets reveals how post-transplantation capabilities are distributed

**DOI:** 10.1101/2024.02.19.579920

**Authors:** Olivia J Stonehouse, Christine Biben, Tom S Weber, Alexandra Garnham, Katie A Fennell, Alison Farley, Antoine F Terreaux, Warren S Alexander, Mark A Dawson, Shalin H Naik, Samir Taoudi

**Affiliations:** The Walter and Eliza Hall Institute of Medical Research, Melbourne, Australia; Department of Medical Biology, The University of Melbourne, Australia; Peter MacCallum Cancer Centre, Melbourne, Australia; Sir Peter MacCallum Department of Oncology, The University of Melbourne, Melbourne, Victoria, Australia; The University of Melbourne Centre for Cancer Research, The University of Melbourne, Melbourne, Victoria, Australia; Lowy Cancer Research Centre, University of New South Wales, Australia; School of Cellular and Molecular Medicine, University of Bristol, Bristol, United Kingdom

## Abstract

It has been proposed that adult haematopoiesis is sustained by multipotent progenitor (MPP) clones that are specified during development. From an immunophenotypic perspective, it is known that hematopoietic stem cell (HSC) and MPPs are present in the fetal liver yet our understanding of how fetal MPPs functionally compare to those in the adult bone marrow is incomplete. Using acute-term transplantations, we found that at a population-level fetal immunophenotypic MPP classes exhibited similar lineage biases as adult cells, albeit with some difference in lymphoid output. Clonal assessment of fetal MPPs engraftment revealed that lineage biases largely resulted from differences in the pattern of single-or bi-lineage differentiation. Immunophenotypic long-term (LT)-and short-term (ST)-HSCs in the fetal liver were distinguished from MPPs according to propensity for clonal multi-lineage differentiation. We also discovered that a large cohort of long-term repopulating units (LT-RU) were within the immunophenotypic ST-HSC population, a significant portion of these were labelled using *Flt3*-cre. This finding has two implications: (1) use of the CD150+ LT-HSC immunophenotype alone will systematically underestimate the size and diversity of the fetal LT-RU pool; and, (2), given fetal LT-RUs with a ST-HSC immunophenotype have the functional attributes required to persist into adulthood.

## Introduction

Hematopoiesis ensures the continuous supply of mature blood lineages. Whether the adult hematopoietic hierarchy is stem cell-driven or is a process sustained by multipotent progenitor cells (MPPs) but underwritten by hematopoietic stem cells (HSCs) is a contentious issue (Busch et al., 2015; Patel et al., 2022; Pei et al., 2017; Pei et al., 2020; Rodriguez-Fraticelli et al., 2018; Sawai et al., 2016; Sawen et al., 2018; Schoedel et al., 2016; Sheikh et al., 2016; Sun et al., 2014).

Based on the ability to provide durable, high-level, and multilineage reconstitution following transplantation, HSCs are traditionally thought to be at the foundation of the hierarchy. Adult bone marrow (ABM) HSCs consist of subsets that can be classified according to durability of self-renewal and by hematopoietic lineage production (Dykstra et al., 2007; Eaves, 2015; Oguro et al., 2013; Pietras et al., 2015): myeloid-biased αHSCs and lineage balanced βHSCs exhibit extensive self-renewal in serial transplantation assays; γHSCs and δHSCs are both lymphoid-biased and fail to reconstitute hematopoiesis following serial transplantation (Dykstra et al., 2007; Muller-Sieburg et al., 2002). In contemporary studies, the most common convention used to resolve HSCs and MPPs is variations of the lineage marker (LIN)-SCA1+ KIT+ (LSK) and SLAM systems, known as the LSK-SLAM code (Kiel et al., 2005; Pietras et al., 2015). Long-term reconstituting HSCs (LT-HSCs) are recognized as LSK_FLT3-CD150+CD48-cells, and short-term reconstituting HSCs (ST-HSCs) are LSK_FLT3-CD150-CD48-(Kiel et al., 2005; Pietras et al., 2015). Most αHSCs/βHSCs are CD150+, and γHSCs/δHSCs are CD150-(Kent et al., 2009).

Downstream of HSCs are the MPPs. ABM MPPs are capable of acute term multilineage differentiation when transplanted into irradiated adult mice (Adolfsson et al., 2001; Kiel et al., 2005; Oguro et al., 2013; Pietras et al., 2015). Although the number of MPP subsets that can be immunophenotypically distinguished in the ABM is increasing (Mansson et al., 2007; Naik et al., 2013; Oguro et al., 2013; Pietras et al., 2015; Sommerkamp et al., 2021), there are three commonly studied MPPs subclasses (MPP2, MPP3, and MPP4). Largely based on studies in the adult, each of the MPP 2 – 4 subsets are capable of lymphoid, myelo-erythroid, and platelet production but exhibit clear lineage biases (Adolfsson et al., 2001; Adolfsson et al., 2005; Oguro et al., 2013; Pietras et al., 2015): MPP2 (LSK_FLT3-CD150+CD48+) and MPP3 (LSK_FLT3-CD150-CD48+) possess limited lymphoid potential; MPP2 exhibits erythroid and megakaryocyte/platelet bias; MPP3 exhibits granulocytic bias; and, MPP4 (LSK_FLT3+CD150-CD48+) exhibits a lymphoid bias. *In situ* barcoding studies suggest that long-term self-renewing MPPs are the drivers of native hematopoiesis because the barcodes present in mature lineages are found in the MPPs but not in LT-HSCs (Patel et al., 2022; Pei et al., 2017; Sun et al., 2014).

Understanding the biology of the HSPC lineages during embryogenesis is important because LT-HSCs (de Bruijn et al., 2000; Ema and Nakauchi, 2000; Ganuza et al., 2017; Gekas et al., 2005; Kumaravelu et al., 2002; Medvinsky and Dzierzak, 1996; Ottersbach and Dzierzak, 2005; Taoudi et al., 2008) and long-lived MPP clones (Patel et al., 2022) emerge during this period. Although the first long-term repopulating units (LT-RUs, a functional designation rather than an immunophenotypic description) express KIT (Sanchez et al., 1996) and SCA1 (de Bruijn et al., 2002) they lack CD150 expression (McKinney-Freeman et al., 2009). By E14.5, CD150 expression enables a high-frequency enrichment of LT-RUs in the fetal liver (Kim et al., 2006), the majority of these are of the βHSC (lineage balanced) class (Benz et al., 2012). Cells with the ST-HSC immunophenotype (CD150-) are present in the E14.5 FL (Kim et al., 2006; Patel et al., 2022), but their functional capacity has not been thoroughly investigated. Although is it known that long-term hematopoietic reconstitution is possible from CD150-cells in the E14.5 FL (Kent et al., 2009; Kim et al., 2006; Papathanasiou et al., 2009), how much of the LT-RU biomass they represent is not known. This is an important issue that requires resolution because of how ubiquitously CD150 expression is used as a hurdle criterion for the quantitative and qualitative analysis of the fetal LT-RU pool (for examples see (Che et al., 2022; Ganuza et al., 2022; Gao et al., 2022; Hall et al., 2022; Lee et al., 2022; Patel et al., 2022; Sakai et al., 2022; Van Deren et al., 2022; Young et al., 2021)).

It has been proposed that the first phase of HSC-derived blood production could be contributed to by developmentally restricted HSCs (drHSCs) (Beaudin et al., 2016). drHSCs derive along a *Flt3-Cre*-expressing ancestry, express the conventional LT-HSC immunophenotype (are LSK CD150+), and provide durable but strongly lymphoid-biased reconstitution following transplantation. drHSCs differ from *bona fide* LT-HSCs according to their ancestry, and the transient nature of their hematopoietic contribution under physiological developmental conditions (Beaudin et al., 2016). One interpretation is that the drHSC subset of the fetal liver LT-HSCs could represent the prenatal equivalent of ST-HSCs (Patel et al., 2022).

MPP4-like features such as transcriptional lineage priming and lymphoid-bias have been reported in the E14.5 fetal liver (Benz et al., 2012; Kim et al., 2006; Mansson et al., 2007). Functional testing of the ABM MPP2 immunophenotype in cells from the E16.5 and E18.5 FL showed evidence of transient multilineage reconstitution (Hall et al., 2022). How functionally comparable the FL and ABM MPP immunophenotypes are remains incompletely understood.

To better understand the functional capability of fetal hematopoietic stem and progenitor cells, we used single cell RNA-Sequencing (scRNA-Seq), cellular barcoding, and transplantation experiments to investigate the E14.5 fetal liver LSK subsets. We found that functionally diverse MPP subtypes do exist within the embryo, and that the adult MPP markers broadly enrich for the same functional groups in the fetus. Clonal tracking following transplantation into adult mice revealed that although fetal MPPs were capable of multi-lineage outcomes in the majority of cases MPPs underwent single-or dual-lineage hematopoietic contribution. Most intriguingly, we discovered that two variants of *bona fide* LT-RUs (without overt linage bias) co-exist in the fetal liver: one is within the LT-HSC immunophenotype and the other within the ST-HSC immunophenotype. Although the LT-RU biomass is distributed between both HSC immunophenotypes, the largest fraction is contained within the ST-HSC immunophenotype. Using *Flt3*-cre lineage tracking we found that ST-HSC immunophenotype was more significantly labelled than LT-HSCs. This suggests that multiple LT-RU forming pathways exist to establish the fetal HSC pool. Consequently, any strategy based on enriching LT-RUs without including those within the CD150-negative fraction will not only omit a large fraction of LT-RUs, but could be missing an important piece of the biological landscape of stem cell pool.

In combination with the insight that the FL ST-HSC population is developmentally distinguished from LT-HSCs according to *Flt3*-expressing ancestry and that embryonic *Flt3*-cre labelled cells contribute to native adult hematopoiesis (Patel et al., 2022), our findings suggest that LT-RUs present in the FL ST-HSC population could persist into adulthood and contribute to native hematopoiesis.

## Materials and Methods

### Mice

Ly5.1, Ly5.1 x C57BL/6, *Rosa26_*floxed-EYFP (Srinivas et al., 2001), UBC-GFP (Schaefer et al., 2001), and *Flt3*-cre (Benz et al., 2012) mice were maintained on a C57BL/6 background. For timed matings, noon of the day of a positive vaginal plug check was considered embryonic day (E) 0.5. Animal experiments were approved by the WEHI animal ethics committee.

### Fetal tissue collection

The uterus/extra-embryonic tissues were washed with Ca^2+^/Mg^2+^-containing DPBS (supplemented with 7% fetal calf serum, 100 units/ml penicillin-streptomycin). Embryos were dissected using watchmaker’s forceps under a microscope (Leica M80) and staged according to morphological features according to Theiler’s criteria. Fetal livers were dissected, mechanically dissociated in 1mL of Ca^2+^/Mg^2+^-free DPBS (supplemented with 7% fetal calf serum, 100 units/ml penicillin-streptomycin, 0.25mM EDTA), and filtered through a 70 μm sieve to prepare single cell suspensions. The total number of cells in each sample was determined using a hemocytometer (BLAUBRAND, Neubauer).

### Adult tissue collection

Femurs and tibias were dissected, washed, and crushed in Ca^2+^/Mg^2+^-free DPBS (supplemented with 7% fetal calf serum, 100 units/ml penicillin-streptomycin, 0.25mM EDTA) using a mortar and a pestle. Bone marrow suspensions were mechanically dissociated, and filtered through a 70 μm sieve. Spleens and thymi were isolated and a single cell suspension were created by massaging each organ through a 70 μm sieve. The total number of cells in each sample was determined using a Neubauer hemocytometer. Blood samples were collected via mandible or retro-orbital bleeds in EDTA-containing microvette tubes.

### Flow cytometry/Cell sorting

KIT-expressing fetal liver or adult bone marrow cells were enriched using magnetic separation (Miltenyi Biotech) according to the manufacturer’s instructions. KIT-enriched cells were stained for flow analysis or sorting using the following immunophenotypes: LT-HSCs were defined as Lin-KIT+SCA1+FLT3-CD48-CD150+; ST-HSCs Lin-KIT+SCA1+FLT3-CD48-CD150-; MPP4s Lin-KIT+SCA1+FLT3+CD48+CD150-; MPP3s Lin-KIT+SCA1+FLT3-CD48+CD150-; MPP2s Lin-KIT+SCA1+FLT3-CD48+CD150+. Lineage cocktail contained; Ter119, B220, MAC1, CD3ε+, CD19, CD4, CD8, Ly6G, NK1.1. Anti-Mac1 antibody was omitted in the lineage cocktail for fetal liver preparations to avoid excluding some of the Lin-KIT+SCA1+ ^(Morrison et al., 1995)^. Differentiated hematopoietic lineages were defined as: erythroid cells TER119+; B cells TER119-MAC1-B220+ CD19+; Monocytes TER119-MAC1+ Ly6G-SSC_A low; Neutrophils TER119-MAC1+ Ly6G+; Platelets TER119-CD41+; Natural killer cells TER119-MAC1-B220-CD3ε-NK1.1+; T cells TER119-MAC1-B220-NK1.1-CD3ε+. All antibody stains were performed on ice in the dark. Analysis or cell sorting were carried out on a BD LSRFortessa or a BD FACSAria (using an 85 μm nozzle), respectively. Dead cells were excluded using 7AAD. Samples were further analyzed using FlowJo software. Antibody details can be found in Table S1.

### Hematopoietic reconstitution assays

Recipient mice were 8 – 10 weeks old Ly5.1 females. Lethal irradiation was delivered as a 1100 rads split dose (2 x 550 rads, three hour gap). Lethally irradiated recipients received 2 x 10^5^ bone marrow filler cells (from Ly5.1 or Ly5.1/2 mice). Sub-lethal irradiation was delivered as a 600 rads dose (no filler cells were used). Short-term (four weeks) and long-term (≥ 16 weeks) transplantations were performed using lethal irradiation. Acute-term transplantations (two weeks) were performed using a sublethal irradiation dose. For transplantation, cells were collected from the bone marrow of adult UBC-GFP mice, prepared in DPBS with 1% FCS and 100 units/ml penicillin-streptomycin, and delivered into recipients by intravenous injection. Bone marrow from UBC-GFP mice were used as donor cells. Cells were injected into recipients approximately three hours after irradiation. Recipients were maintained on neomycin for 14 – 21 days. For estimation of LT-RU frequency (limiting dilution analysis), reconstitution frequency from the transplantation of 10 – 100 purified cells was analyzed using the extreme limiting dilution method (Hu and Smyth, 2009).

### Single cell RNA-Seq

Cells were index-sorted on a 384 well plate using an BD FACSAriaIII or Fusion. 9 C57BL/6 E14.5 fetal livers were pooled from two litters and sorted into a 384-well plates. Libraries were prepared using CELSeq2 (Hashimshony et al., 2016) with additional optimizations, as described in (Amann-Zalcenstein et al., 2020). E14.5 fetal livers were pooled from two litters and sorted on the same day across multiple replicate plates. Sorts were undertaken using aseptic protocols and using a 100 μm nozzle. For flow alignment, 1 bead coated with Horse Radish Peroxidase was sorted into 1.2 μL of horse-radish peroxidase substrate before and after sort controls. Colorimetric assessment confirmed proper alignment of the cell sorter. Single cells were sorted into plates containing a primer/lysis mix (dNTPs, ERCC and SUPERase Inhibitor). Plates were centrifuged for 1 min at 1,200 x *g* and immediately frozen down at –80°C until further processing. For sequencing, libraries from a single plate (containing 375 single cells) were prepared with a CELSeq2 protocol (Amann-Zalcenstein et al., 2020). Sequencing reads were aligned and mapped to the mm10 mouse genome and ERCC spike-in sequences using Rsubread (Liao et al., 2019). GRCm38.88_chr annotation was then added. The data was demultiplexed and reads overlapping genes were summarized into counts using scPipe v1.6.0 (Tian et al., 2018). All subsequent analyses were performed in R version 3.6.0. Seven cells were identified as outliers based on number of genes detected, total gene counts, ERCC percentage, mitochondrial gene percentage and ribosomal gene percentage, and were excluded leaving 368 cells for downstream analysis. Gender-related genes were removed to avoid potential gender biases in the analysis. Lowly expressed genes were also filtered out such that 15590 genes showing an average count greater than 1 in at least 10 cells were retained. Normalisation factors were computed by deconvolution using scran v1.12.1 (Lun et al., 2016b) and subsequently used to calculate log-transformed normalised expression values. Gene-specific variance of the biological and technical components of the data were then estimated. Cell cycle phases were predicted using the cyclone function in scran. Differential expression analyses both between cell types and between cell clusters were carried out using edgeR v3.26.6 (Chen et al., 2016b; Lun et al., 2016a). For each analysis, biological variation between samples were estimated using edgeR’s estimateDisp function (Chen et al., 2016b). Generalized log-linear models were then fitted to the count data, incorporating an adjustment for cell cycle phase. edgeR’s likelihood ratio test pipeline was applied to identify differentially expressed genes between the cell populations (McCarthy et al., 2012). The false discovery rate (FDR) was controlled below 5% using the Benjamini Hochberg method. Gene Ontology and KEGG pathway analysis were performed using limma v3.40.6 (Ritchie et al., 2015). Dimension reduction using Principal Components Analysis (PCA), Uniform Manifold Approximation and Projection (UMAP), and clustering was performed using Seurat v3.0.2 (Stuart et al., 2019). Code used for analysis will be made available on request.

### Lentiviral barcoding

The mCherry-expressing SPLINTR library of DNA barcodes used contained ∼70,000 unique barcodes (Fennell et al., 2022). E14.5 LSK subsets were collected from pooled littermates (*n* = 4 independent experiments). A maximum of 3 x 10^4^ purified E14.5 FL LSK cells were transduced with SPLINTR virus in 96-well plates via spinfection (90 mins at 1250 x *g*) at a multiplicity of infection of 0.01 - 0.02 (this provided 1 – 2% mCHERRY+ cells) to limit multiple integrations of the barcode per cell. Cells were washed and then injected intravenously into recipient mice. After two weeks mCHERRY+ barcoded cells were retrieved from the spleens of recipient mice two-weeks post transplantation. Spleens were stained with markers for the isolation of monocytes (TER119^-^ MAC1^+^Ly6G^-^ [low side-scatter]), neutrophils (TER119^-^MAC1^+^LY6G^+^), erythroblasts (TER119^+^CD71^+^), and B cells (TER119^-^MAC1^-^B220/CD19^+^). mCHERRY^+^ cell populations were sorted and spun into pellets of no more than 5 x 10^5^ cells. Pellets were resuspended in 40 μL Direct PCR Lysis Reagent (Cell) with 0.5 mg/mL Proteinase K. Cells underwent lysis at 55°C for 2 hrs, followed by 85°C for 30 mins and 95°C for 5 mins. Libraries were generated as outline in (Naik et al., 2013). Raw sequencing FASTQ files were processed using custom C++ code to count SPLINTR barcode reads in each sample, including exclusively barcodes that aligned to a previously generated reference library. All subsequent analyses were performed using the statistical computing language R. Counts below a threshold of 100 reads were set to zero. Barcodes below a threshold of 100 sequencing reads, or only detected in one PCR replicate were removed to minimise artefacts (Figure S1). Heatmaps of the barcode counts were generated using the R package “heatmap 3”. Finally, the fate (presence/absence in B/Ery/Mono/Neut) for each barcode was determined, and correlation between distribution of fates computed between different populations (iLT-HSC/iST-HSC/iMPP2/iMPP3/iMPP4) using the Pearson correlation coefficient. To quantify the contribution of each ancestral clone to a given lineage (lineage biomass), the proportion of reads attributed to each fate combination was computed per LSK subset.

### General statistical analysis

Prism 9 (GraphPad) was used for data analysis and graph production. Data are represented as mean ± standard deviation (SD), and analyzed using Student’s t-test (two-way, unpaired). One-way ANOVA (using Sidak’s P value adjustment) was used for multiple comparisons. ‘*n’* was used to designate the number of independent experimental mice. ns, not statistically significant. * = p<0.05, ** = p<0.01, *** = p<0.001, **** = p<0.0001.

## Results

### Quantitative and transcriptional investigation of fetal LSK subsets

We used the adult LSK-SLAM immunophenotypes (Pietras et al., 2015) to investigate the development of HSPC (LSK) subsets in E11.5 - E14.5 fetal livers (Figure 1A – C). To avoid the presumption of functional equivalence to adult bone marrow HSPCs, we adopted the convention of referring to cell populations with the immunophenotypic (‘i’) prefix (e.g. iST-HSC), as has been implemented previously by others (Chen et al., 2016a; Dong et al., 2020). At E11.5 iST-HSC (LSK_FLT3-CD150-CD48-), iMPP3 (LSK_FLT3-CD150-CD48+), and iMPP4 (LSK_FLT3+CD150-CD48+) populations were observed in the liver (Figure 1A – E). By E12.5 all LSK subtypes, including iLT-HSCs (LSK_FLT3-CD150+CD48-) and MPP2 (LSK_FLT3-CD150+CD48+), were detected (Figure 1A – E).

**Figure 1.**
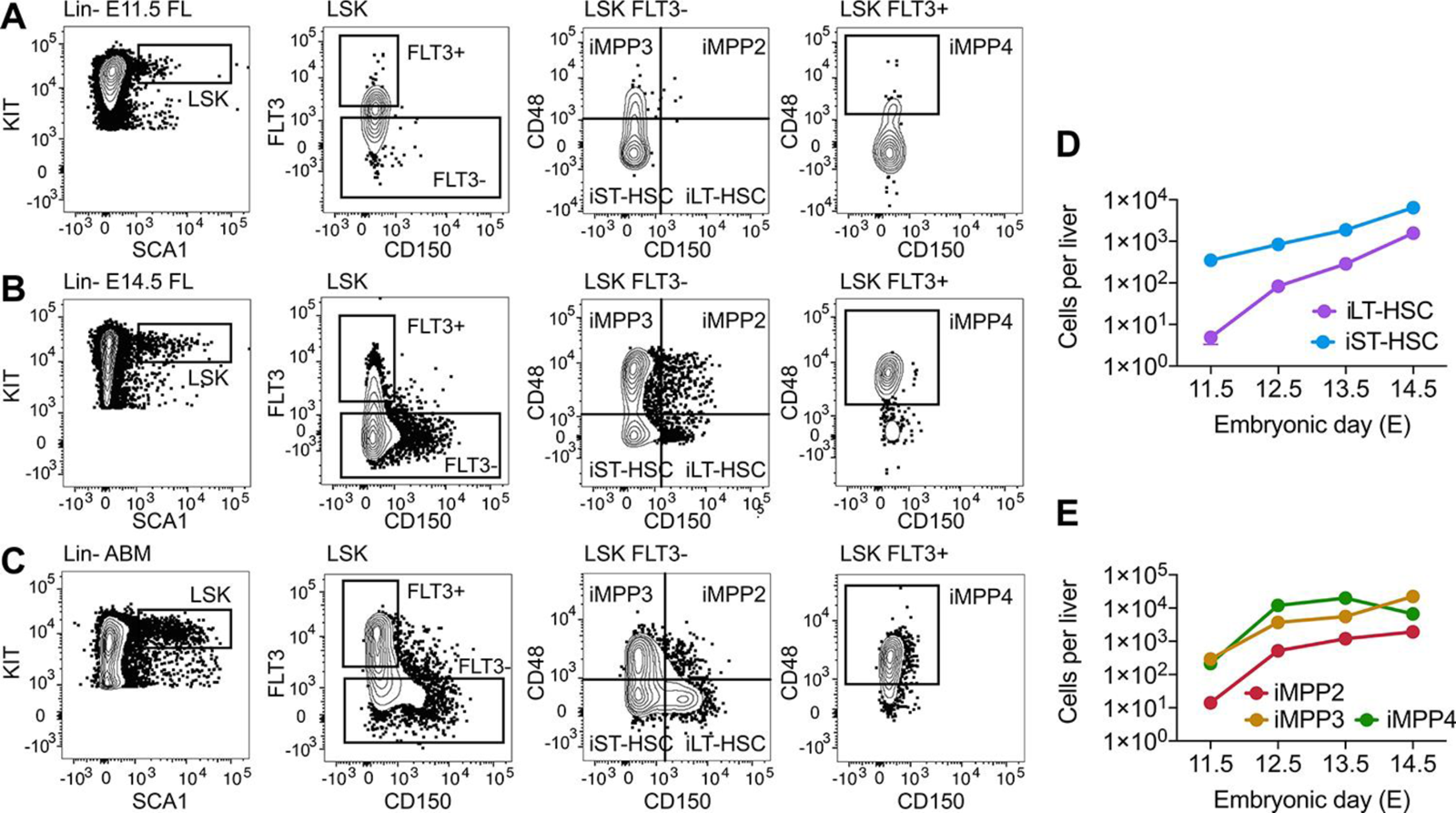
Quantification of LSK subsets in the fetal liver between E11.5 – E14.5. **(A – C)** Representative flow cytometry plots showing the identification of LSK subsets in the E11.5 fetal liver (FL) **(A)**, E14.5 fetal liver **(B)** and adult bone marrow **(C)**. **(D)** Quantification iLT- and iST-HSCs in the fetal liver between E11.5 – E14.5. E11.5 *n* = 7 individual embryos; E12.5, E13.5, and E14.5 *n* = 6 individual embryos per developmental stage. **(E)** Quantification of iMPP2, iMPP3, and iMPP4 in the fetal liver between E11.5 – E14.5. E11.5 *n* = 7 individual embryos; E12.5, E13.5, and E14.5 *n* = 6 individual embryos per developmental stage. iMPP2, immunophenotypic multipotent progenitor 2. iMPP3, immunophenotypic multipotent progenitor 3. iMPP4, immunophenotypic multipotent progenitor 4. iLT-HSC, immunophenotypic long-term hematopoietic stem cell (iLT-HSC). iST-HSC, immunophenotypic short-term hematopoietic stem cell.

We next compared the transcriptomes of E14.5 FL LSK subsets to gain insight into potential functional differences. To this end, single cells from E14.5 FL LSK subsets were purified (as indicated in Figure 2A) using indexed flow cytometry (which recorded the immunophenotype of the purified cell). Index data was used to verify the immunophenotype of the purified cells. Unsupervised hierarchical clustering of single-cell RNA-Sequencing data indicated that E14.5 FL iMPP 2-4 were transcriptionally distinct subsets (Figure 2B). Gene ontology (GO) term analysis using significantly differentially expressed genes between iMPP 2 – 4 revealed:

- Significant enrichment of pathways related to lymphoid differentiation/function in iMPP4s compared to iMPP2 and iMPP3 (Figures 2Ci and 2Di; Tables S2 and S3). This was in keeping with the lymphoid bias of ABM iMPP4 (Pietras et al., 2015).
- Significant enrichment of pathways associated with megakaryocytes and erythroid differentiation in iMPP2 compared to iMPP3 and iMPP4 (Figures 2Di and 2Ei; Tables S3 and S4). This was consistent with the megakaryocyte/erythroid bias of ABM iMPP2.
- Interestingly, in comparison with iMPP2 and iMPP4, iMPP3 presented with a more lineage-balanced transcriptional priming that spanned the erythro-myeloid, megakaryocytic, and lymphoid lineages (Figures 2Ci and 2Ei; Tables S2 and S4).

**Figure 2.**
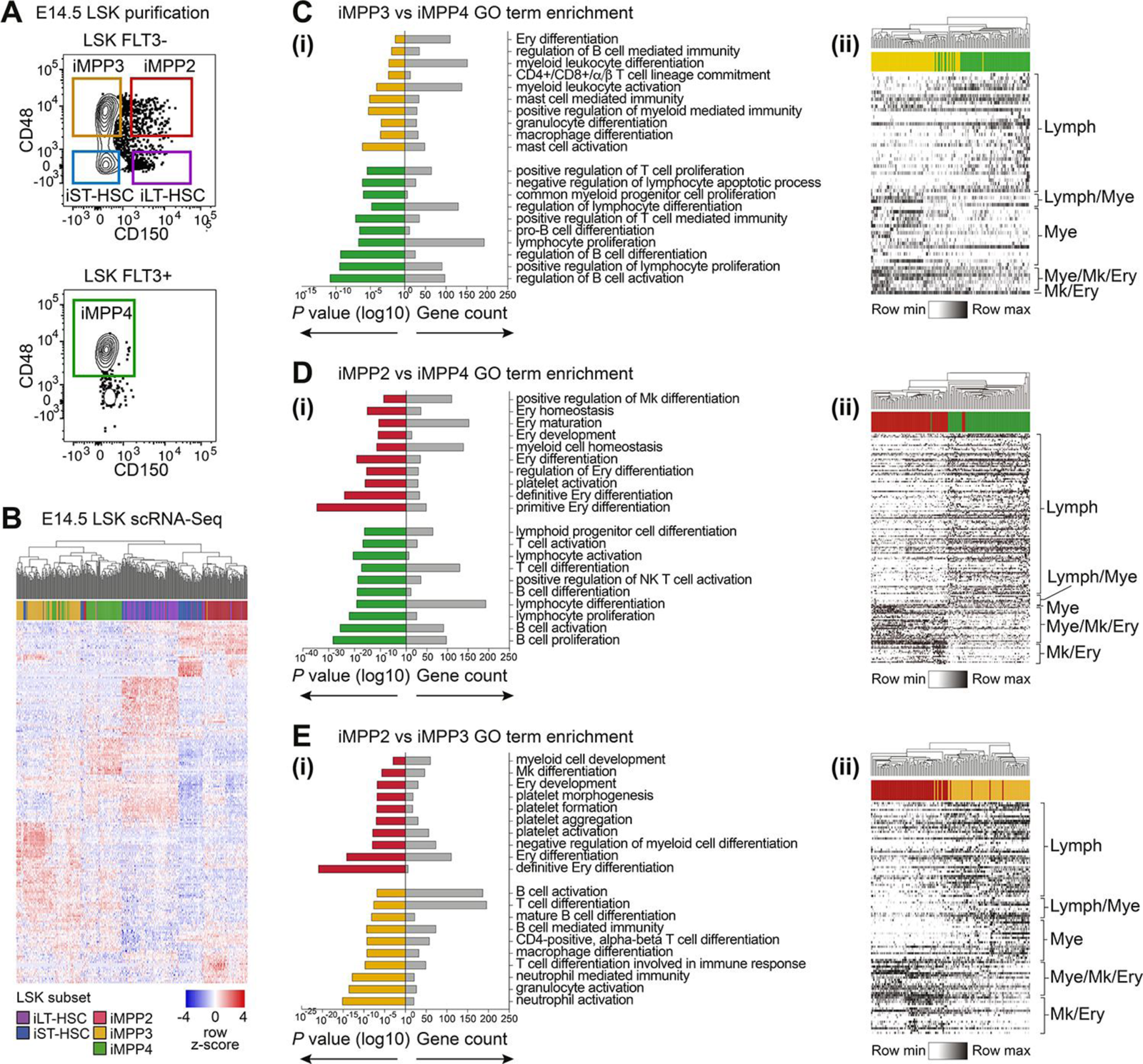
Transcriptional investigation of iMPP subsets in the E14.5 fetal liver. **(A)** Representative example of the gating strategy used to purify LSK subsets from the E14.5 fetal liver. **(B)** Heatmap of the top 100 most variable genes for each E14.5 FL LSK subtype. **(C)** Gene Ontogeny (GO) term enrichment analysis for significantly differentially expressed genes for iMPP3 versus iMPP4 **(i)**. **(ii)** Heatmap of genes present in significantly enriched GO terms. **(D)** Gene Ontogeny (GO) term enrichment analysis for significantly differentially expressed genes for iMPP2 versus iMPP4 **(i)**. **(ii)** Heatmap of genes present in significantly enriched GO terms. **(E)** Gene Ontogeny (GO) term enrichment analysis for significantly differentially expressed genes for iMPP2 versus iMPP3 **(i)**. **(ii)** Heatmap of genes present in significantly enriched GO terms. Fetal liver (FL). Lineage-KIT+SCA1+ (LSK). iMPP2, immunophenotypic multipotent progenitor 2. iMPP3, immunophenotypic multipotent progenitor 3. iMPP4, immunophenotypic multipotent progenitor 4. iLT-HSC, immunophenotypic long-term hematopoietic stem cell (iLT-HSC). iST-HSC, immunophenotypic short-term hematopoietic stem cell. Lymphoid (Lymph). Lymphoid and myeloid (Lymph/Mye). Mye (Myeloid). Mk/Ery (Megakaryocyte and erythroid). Myeloid, megakaryocyte and erythroid (Mye/Mk/Ery).

Visualization of the genes associated with enriched GO terms confirmed that the associated genes were broadly expressed within the relevant populations (Figures 2Cii – Eii and Table S5). This indicated that adult MPP immunophenotypes effectively enriched for E14.5 LSKs with features of ABM-like lineage priming.

### Functional comparison of E14.5 FL and ABM immunophenotypic counterparts in vivo

To compare the acute-term *in vivo* differentiation potential of E14.5 FL and ABM LSK subtypes, cells purified from UBC-GFP mice (Schaefer et al., 2001) were transplanted into GFP-sub-lethally irradiated recipients. To enable a like-for-like comparison the number of cells transplanted was determined by the availability of cells in the fetal liver, and the number of cells that would yield robust engraftment. From either E14.5 FL or ABM donors, 1200 iLT-HSCs, 4000 iST-HSCs, 7000 iMPP2s, 25000 iMPP3s, and 30000 iMPP4s were transplanted. Analysis of *in vivo* output was performed after two weeks. This time point was selected to ensure that the output of iMPPs (which rapidly exhaust (Pietras et al., 2015)) would be captured. Contribution to platelets, erythrocytes, myeloid, and lymphoid lineages was assessed in the peripheral blood, spleen, or bone marrow of recipient mice.

Inter-developmental stage comparison of the HSC subsets revealed largely equivalent outcomes (Figure 3A and B), the exception was greater erythroid output by FL iLT-HSCs (Figure 3A). Of note, the low T-cell observed from all of the HSC subsets two weeks after transplantation was consistent with observations from other studies (Forsberg et al., 2006; Papathanasiou et al., 2009; Pietras et al., 2015). iMPP 2 – 4 subsets also performed similarly between the developmental stages (Figure 3 C – E). The exceptions were enhanced B-cell output from FL iMPP3 (Figure 3D) and diminished T-cell output from FL iMPP4 (Figure 3E).

**Figure 3.**
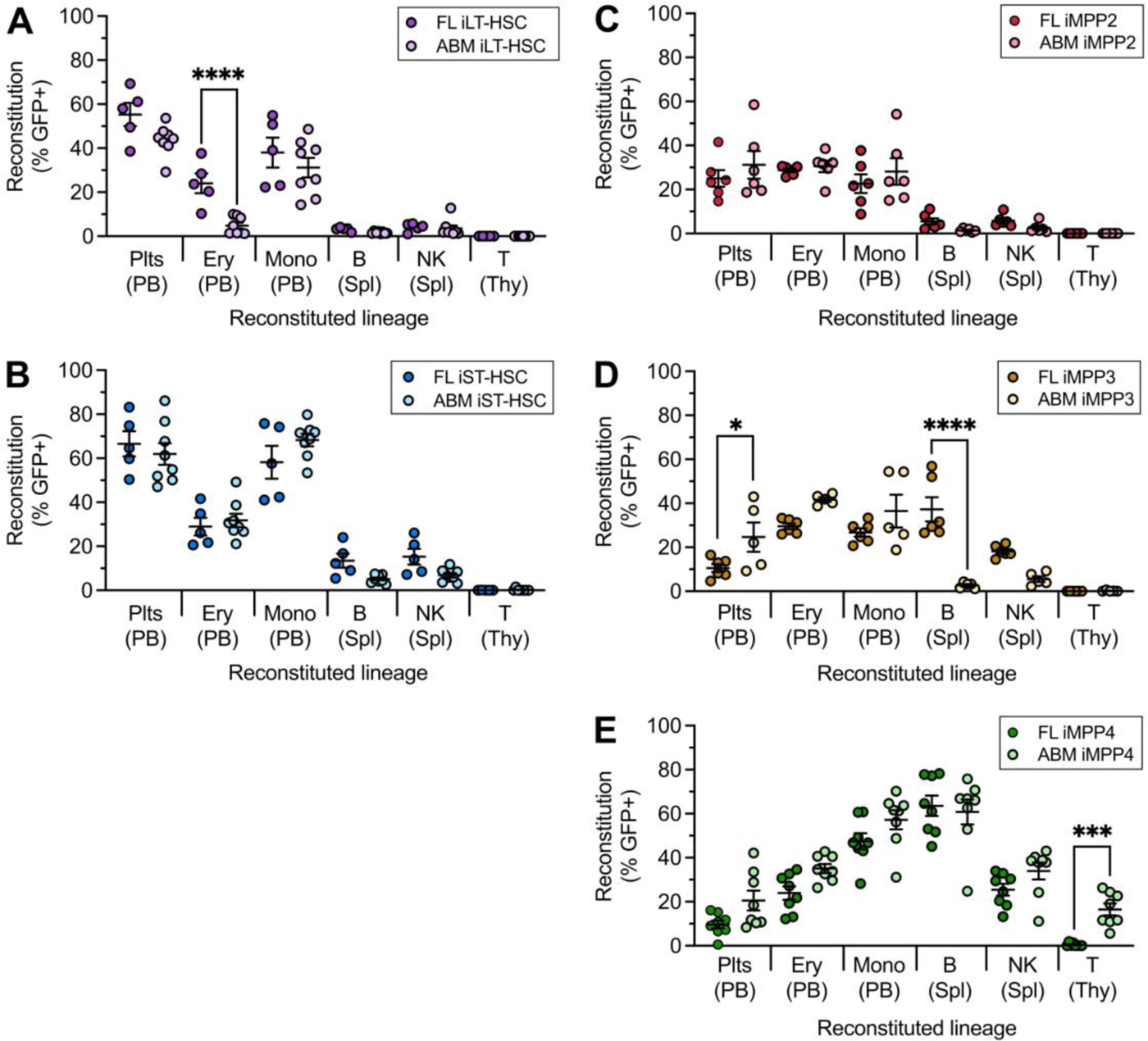
Functional comparison of E14.5 FL LSK subsets with ABM counterparts in acute-term hematopoietic reconstitution assays. Comparison of multi-lineage reconstitution between E14.5 FL and ABM iLT-HSCs (**A**), iST-HSCs (**B**), iMPP2 (**C**), iMMP3 (**D**), and iMPP4 (**E**) two weeks after transplantation. Data were analyzed using one-way ANOVA. The Šídák test was used to correct for multiple comparisons. ns, not statistically significant. *, *p* < 0.05. **, *p* < 0.005, ***, *p* < 0.0005, ****, *p* < 0.00005. ABM iLT-HSC, *n* = 8 independent recipient mice. ABM iST-HSC, *n* = 8 independent recipient mice. E14.5 iLT-HSC, *n* = 4 independent recipient mice. E14.5 iST-HSC *n* = 5 independent recipient mice. ABM iMPP2, *n* = 6 independent recipient mice. ABM iMPP3, *n* = 5 independent recipient mice. ABM iMPP4, *n* = 8 independent recipient mice. E14.5 iMPP2, *n* = 6 independent recipient mice. E14.5 iMPP3, *n* = 6 independent recipient mice. E14.5 iMPP4, *n* = 7 independent recipient mice. iMPP2, immunophenotypic multipotent progenitor 2. iMPP3, immunophenotypic multipotent progenitor 3. iMPP4, immunophenotypic multipotent progenitor 4. iLT-HSC, immunophenotypic long-term hematopoietic stem cell (iLT-HSC). iST-HSC, immunophenotypic short-term hematopoietic stem cell. Plt, Platelet. Ery, erythroid. Mono, Monocyte. B, B-cell. T, T-cell. NK, natural killer cell. Parentheses, analyzed tissue. PB, peripheral blood. Spl, spleen. Thy, thymus.

### Clonal tracking E14.5 FL LSK subsets following transplantation

To understand the clonal nature of *in vivo* differentiation of FL iHSC and iMPP subsets we transplanted cells following lentiviral cellular barcoding. *Ex vivo* cellular barcoding involves the indelible labelling of individual cells with a high-diversity library of genetically heritable DNA sequences termed barcodes (Fennell et al., 2022; Gerrits et al., 2010; Lu et al., 2011; Naik et al., 2013; Schepers et al., 2008). Once LSK subsets are transduced with a barcode-carrying lentivirus they can be transplanted into irradiated recipients and their barcoded progeny can be purified by flow cytometry. Clonal ancestry can be determined by retrieving barcode sequences from the genome of donor-derived cells.

Purified FL LSK subsets were transduced with a previously described mCHERRY-expressing lentiviral barcode library composed of ∼ 7 x 10^4^ unique barcodes (Fennell et al., 2022). A maximum of 3 x 10^4^ cells were cultured with a pre-titered library for 16 hrs to achieve 1 – 2 % transduction. This provided a >100-fold excess in barcode diversity, making repeat use of barcodes and multiple infection of individual cells unlikely (Kebschull and Zador, 2018; Naik et al., 2014) (Figure S1). Of note, the culture period did not affect *in vivo* output (Figure S2). Two weeks after transplantation, mCHERRY+ B-cells, erythroblasts, monocytes, and granulocytes were harvested from recipient mice. It has been previously shown that at two weeks post-transplantation the spleen not only contains the offspring of clones that also engraft the bone marrow but it also contains clones that have not engrafted the bone marrow (Naik et al., 2013). Accordingly, to ensure effective retrieval of barcoded offspring we focused on the retrieval of hematopoietic lineages from the spleen to determine the patterns of clonal outcomes for each of the E14.5 LSK subsets (Figure 4A).

**Figure 4.**
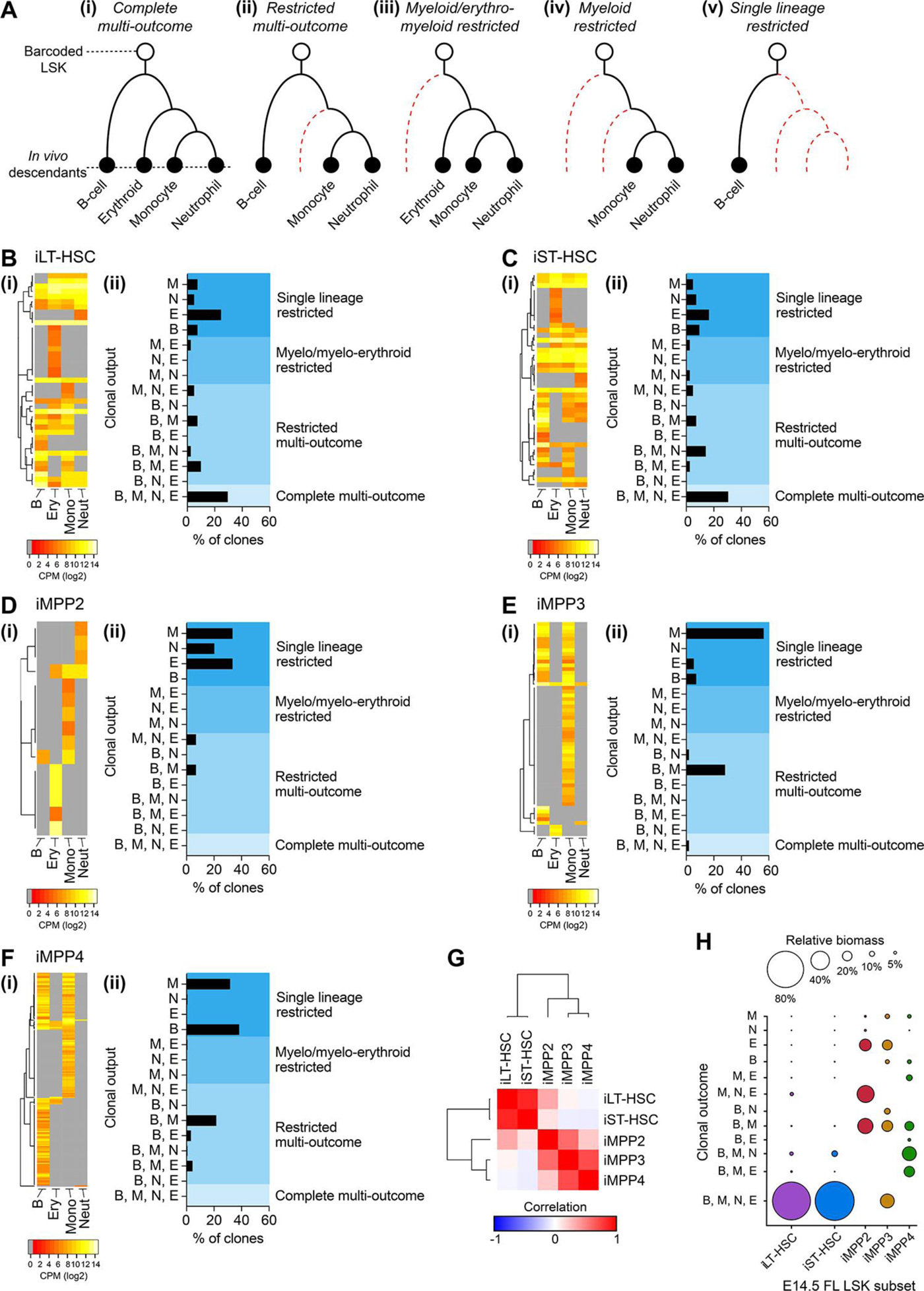
Clonal multilineage outcomes are restricted to the E14.5 iLT-HSCs and iST-HSCs. **(A)** Clonal output from barcoded E14.5 LSK subsets was determined by retrieving barcodes in donor-derived hematopoietic lineages two weeks after transplantation. Based on the lineages with shared barcodes, clones were classified as having either: **(i)** Complete multi-lineage outcome (complete multi-outcome) when the same barcode was retrieved from each lineage; **(ii)** Restricted multi-outcome when contribution to B-cells and to at least one of the erythro-myeloid lineages; **(iii)** Erythro-myeloid restricted when contribution to only erythroblasts, monocytes, and granulocytes was observed; **(iv)** Myeloid restricted when contribution to only monocytes, and granulocytes was observed; **(v)** Single-lineage restricted when contribution to only one of the lineages was observed. **(B-F)** Heatmaps of barcodes reads **(i)** and histograms summarising the frequencies of clonal outputs **(ii)** for iLT-HSC (**B**, *n* = 41 clones), iST-HSC (**C**, *n* = 43 clones), iMPP2 (**D**, *n* = 15 clones), iMPP3 (**E**, *n* = 57 clones), and iMPP4 (**F**, *n* = 168 clones). M, monocyte. N, neutrophil. E, erythroid cell. B, B-cell. CPM, barcode sequence counts per million reads. **(G)** Similarity matrix analysis of global clonal outcomes of E14.5 LSK subsets. **(H)** Bubble plot summary of the proportion of reads associated with specific clonal outcomes (relative biomass) for the E14.5 LSK subtypes. Rows are the clonal outcomes (lineages produced). Size of the bubble represents the relative biomass contribution. M, monocyte. N, neutrophil. E, erythroid cell. B, B-cell.

E14.5 FL iLT-HSC clones (*n* = 41) and FL iST-HSC clones (*n* = 43) contributed to similar clonal outputs (Figure 4B and 4C). This ranged from complete multi-lineage outcomes that spanned the lymphoid, myeloid, and erythroid lineages (∼30% of clones), various combinations of restricted multi-outcomes, erythro-myeloid restriction, and single-lineage restriction (Figure 4B and 4C). This was in stark contrast to the outcome of MPPs. Clonal tracking of MPPs revealed that:

- Of iMPP2 clones (*n* = 15), some were capable of restricted multi-outcomes but 86% of clones underwent single-lineage myeloid or erythroid production (Figure 4D).
- Of iMPP3 clones (*n* = 57), 68% were single-lineage restricted; this included production of B-cells, monocytes, or erythroblasts. 30% of clones underwent restricted multi-outcomes (myeloid lineage and B-cell outcome), and only 1 clone underwent complete multi-outcome (Figure 4E).
- Although similar in the performance to iMPP3, iMPP4 clones (*n* = 168) were more prone to B-cell restricted outcome (Figure 4F).

Although all iMPPs (2 – 4) exhibited some degree of multi-lineage outcomes, contribution to all tested lineages was a feature largely restricted to iLT-HSCs and iST-HSCs.

Correlation between distribution of outcomes between the LSK subsets revealed a marked difference between iLT-HSCs and iST-HSCs compared to the iMPPs (Figure 4G). The iHSC versus iMPP segregation was also evident when contribution of clones to lineage biomass was considered (Figure 4H). These data revealed that E14.5 iLT-HSCs and iST-HSCs contained cells with the capacity for complete multi-lineage outcomes, and that the lineage-biased outcomes observed from MPPs at the population-level were likely driven by a mix of clones that underwent with single-lineage restricted outcomes.

### Comparison of stemness between E14.5 FL iLT-HSCs and iST-HSCs

From the barcoding experiments we noted that FL iLT-HSCs and iST-HSCs performed similarly. ABM iST-HSCs diverged from ABM iLT-HSCs according to differences in durability of multilineage reconstitution, and by the inability to self-renew (Morita et al., 2010; Oguro et al., 2013; Pietras et al., 2015). To further investigate FL iST-HSCs, we next compared the function of E14.5 FL iLT-HSCs and iST-HSCs in long-term transplantation assays. To distinguish immunophenotypic classification from stem cell function we will continue to refer to HSC immunophenotypes as either iLT-HSC or iST-HSC but the provision of durable multilineage reconstitution will be referred to as the output of a long-term repopulating unit (LT-RU).

We first compared the performance of E14.5 FL iLT-HSCs and iST-HSCs following transplantation into lethally irradiated recipients (the gold-standard HSC assay). Analysis of erythroid, platelet, and leucocyte reconstitution at 5, 16, and 40 weeks after transplantation of 1000 cells per recipient indicated that *bona fide* LT-RUs were present in both populations (Figure 5A). Analysis of the LSK compartment of recipients (after 40 weeks) revealed that LT-RUs from both populations generated all ABM LSK subsets, including iLT-HSCs (Figure 5B).

**Figure 5.**
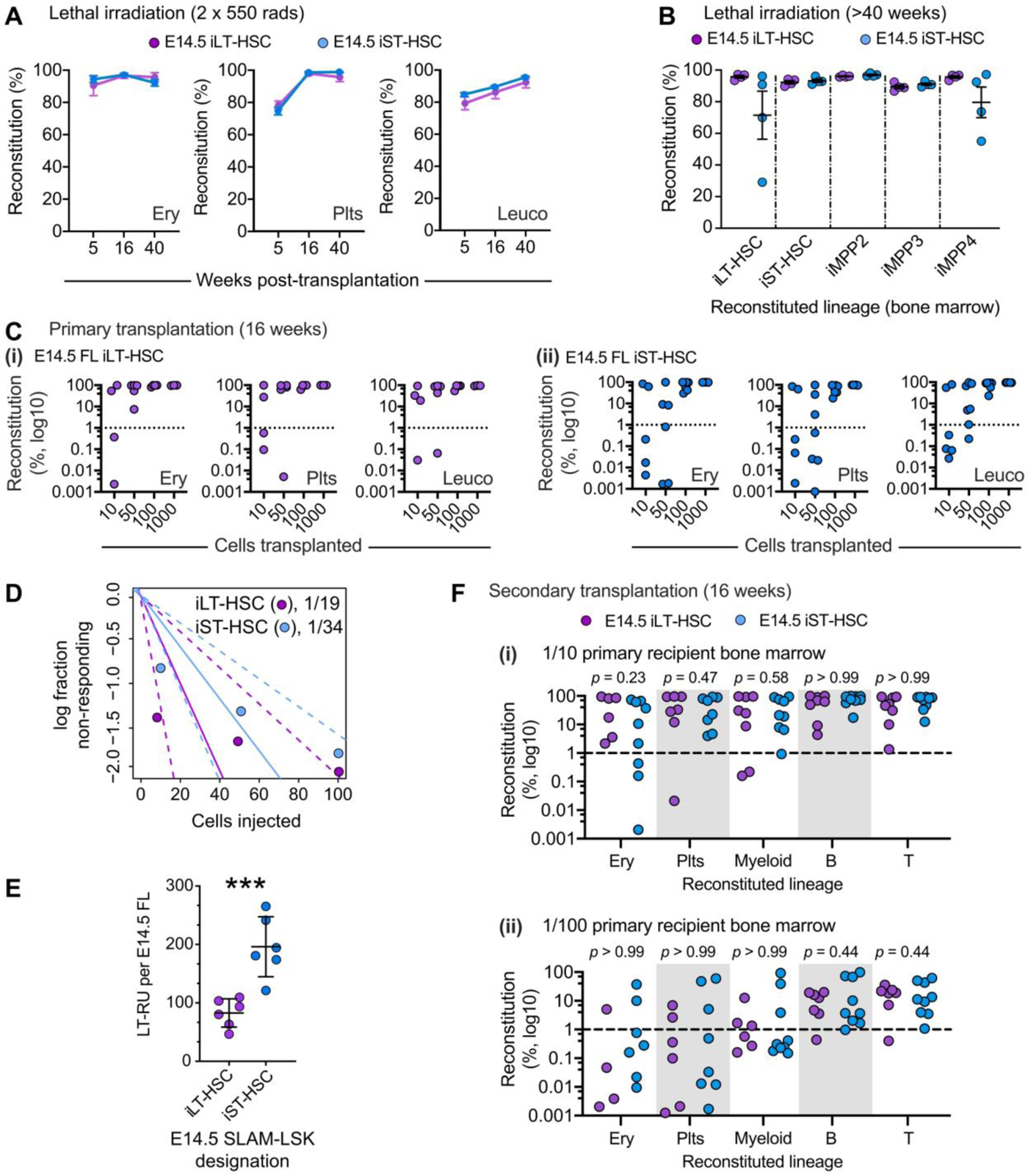
Large numbers of LT-RUs are concealed in the E14.5 iST-HSC immunophenotype. **(A)** 1,000 E14.5 FL iLT-HSCs (purple) or iST-HSCs (blue) were transplanted into lethally irradiated (2 x 550 rads) recipients. Peripheral blood platelets (Plt), erythrocytes (Ery), and leucocytes (leuco) were analyzed at 5, 16, and 40 weeks post transplantation. iLT-HSCs, *n* = 10 recipient mice. iST-HSCs, *n* = 11 recipient mice. **(B)** Reconstitution of ABM LSK subsets following transplantation with 1,000 E14.5 FL iLT-HSCs or iST-HSCs in lethally irradiated recipients >40 weeks after transplantation. iLT-HSCs, *n* = 4 recipient mice. iST-HSCs, *n* = 4 recipient mice. **(C)** Titration of E14.5 iLT-HSCs **(i)** or iST-HSCs **(ii)**. 10, 50, 100 or 1000 cells were injected into lethally irradiated recipients. Peripheral blood platelets (Plt), erythrocytes (Ery), and leucocytes (leuco) were analyzed 16 weeks after transplantation. Dashed line represents the 1% reconstitution threshold. **(D)** Limiting dilution analysis was used to estimate long-term repopulating unit (LT-RU) frequency in the E14.5 iLT-HSC and iST-HSC populations. LT-RU frequency was 1/19 of iLT-HSCs and 1/34 of iST-HSCs. Dotted lines represent 95% confidence interval for each population. **(E)** Estimation of absolute numbers of LT-RUs in the E14.5 iLT-HSC and iST-HSC populations. iLT-HSC *n* = 6 individual embryos. iST-HSC *n* = 6 individual embryos. **(F)** Secondary transplantations were performed using 1/10 **(i)** or 1/100 **(ii)** doses of primary recipients that had received either a 100-cell dose of E14.5 iLT-HSCs or of iST-HSCs. Peripheral blood platelets (Plt), erythrocytes (Ery), and leucocytes (leuco) were analyzed 16 weeks after transplantation. For (i) iLT-HSC secondary recipients, *n* = 6 secondary recipients from 3 independent primary recipients; iST-HSC secondary recipients, *n* = 9 secondary recipients from 3 independent primary recipients. For (ii) iLT-HSC secondary recipients, *n* = 6 secondary recipients from 3 independent primary recipients; iST-HSC secondary recipients, *n* = 8 secondary recipients from 3 independent primary recipients. Dashed line represents the 1% reconstitution threshold. *p* values are derived from contingency analysis (using Fisher’s exact test) of reconstitution outcomes (above versus below the 1% reconstitution threshold) from iLT-HSCs and iST-HSCs for each lineage.

We next performed limiting dilution experiments to determine the frequency of LT-RUs in the FL iLT-HSC and iST-HSC populations. For this, we transplanted 10, 50, 100, or 1000 purified cells into lethally irradiated recipients; mice were analyzed after 16 weeks (Figure 5C). Using the extreme limiting dilution method (Hu and Smyth, 2009), we found that LT-RUs were present at a frequency of 1/19 iLT-HSCs (which is in keeping with a previous estimate (Kim et al., 2006)) and 1/34 iST-HSCs (Figure 5D). Considering the absolute number of iLT-HSCs and iST-HSCs per E14.5 FL (Figure 1B), we estimated that the iST-HSC population contained approximately twice the number of LT-RUs than the iLT-HSC population (Figure 5E).

To investigate the self-renewal capacity of LT-RUs in the iST-HSC population (herein referred to as iST:LT-RU, 1/10 or 1/100 doses of ABM from primary recipients (that had received 100 donor cells) were transplanted into lethally irradiated secondary recipients. We found no significant difference in secondary reconstitution between iLT:LT-RU and iST:LT-RU (Figure 5F). Thus, the E14.5 FL ST-HSC population contains large number of *bona fide* LT-RUs that did not exhibit notable lineage biases.

### Refinement of the E14.5 FL ST-HSC LT-RU immunophenotype

We next investigated our E14.5 FL LSK scRNA-Seq dataset to understand if a unifying LT-RU transcriptional signature could be identified. Based on similarity matrix analysis, 7 transcriptional clusters were identified; most iLT-HSCs and iST-HSCs segregated into clusters 4 and 7 (Figure 6A). Derivation of signature genes for these clusters indicated that cluster 4 likely contained LT-RUs: *Mecom*, *Hlf*, and *Mllt3* are strongly associated with LT-RU function (Calvanese et al., 2019; Kataoka et al., 2011; Komorowska et al., 2017; Wahlestedt et al., 2017) and were enriched in cluster 4 (Figure 6A, Figure S3, Table S6, and Table S7). In contrast, erythroid-associated genes (e.g. *Epor*, *Klf1*, and *Redrum*) were expressed at higher level in cluster 7 (Table S6 and Table S7). Intra-cluster differential gene expression analysis between within iLT-HSCs and iST-HSCs revealed that cells within clusters were transcriptionally very similar (Figure 6B, Table S8, and Table S9), while many genes were differentially expressed between clusters (Figures 6C, Table S6, and Table S7). Although very similar, five genes were differentially expressed between iLT-HSCs and iST-HSCs within cluster 4: *Gm42418*, *Igf2*, *Igf1r*, *Mir6236*, and *Satb1* were more highly expressed in iST-HSCs (Table S8). Previous studies have shown that: *in vitro* treatment of FL and ABM iLT-HSCs with IGF2 results in LT-RU expansion (Zhang and Lodish, 2004); level of IGF2 receptor expression in FL and ABM cells correlates with greater hematopoietic reconstitution (Zhang and Lodish, 2004); enforced expression of *Igf2* in ABM iLT-HSCs enhances hematopoietic reconstitution (Thomas et al., 2016); and, high-levels of *Satb1* expression correlates with improved hematopoietic reconstitution (Doi et al., 2018). This suggests that some degree of functional difference between iLT:LT-RUs and iST:LT-RUs might exist.

**Figure 6.**
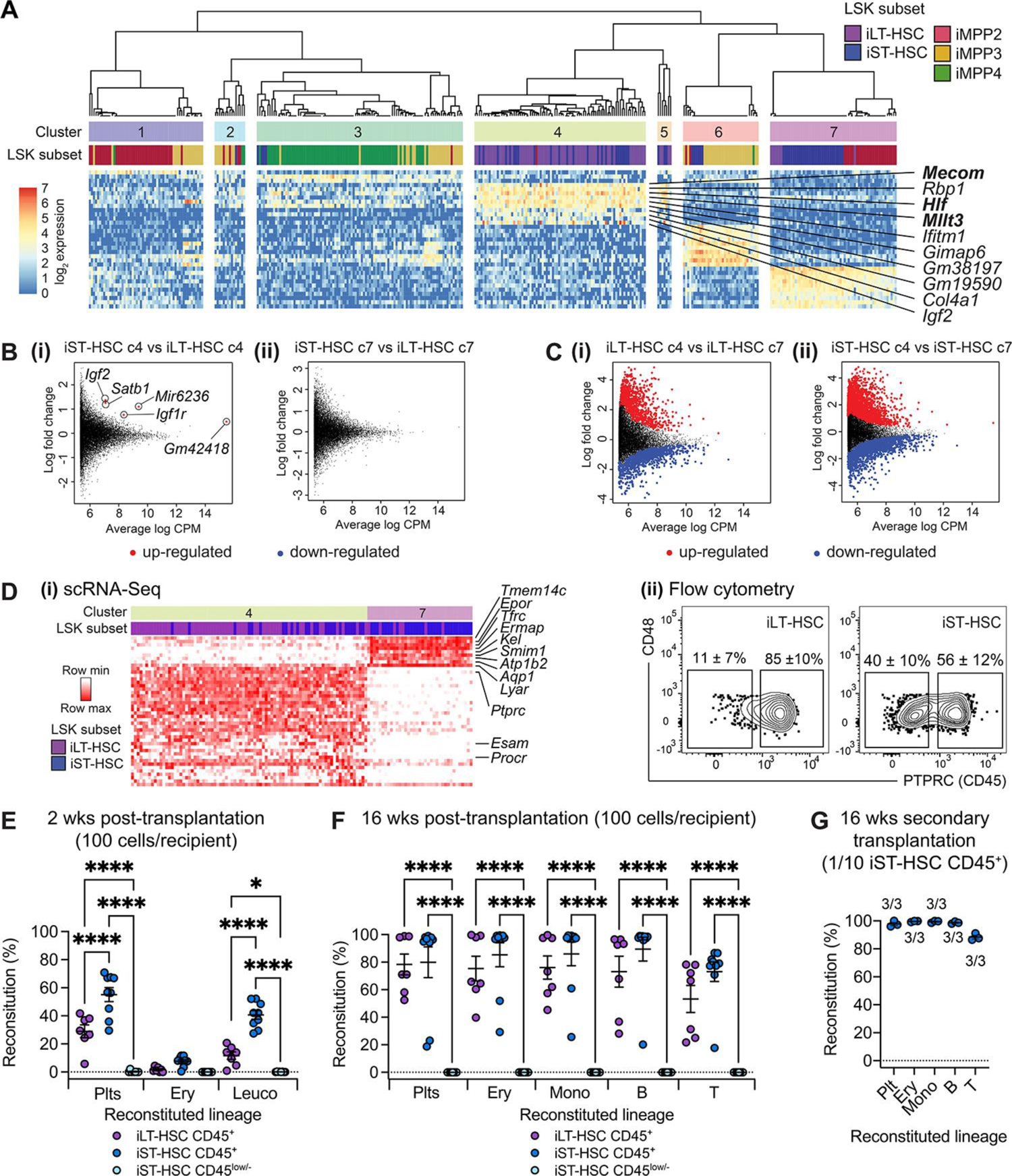
Use of single-cell transcriptomics to define a more refined iST:LT-RU immunophenotype. **(A)** E14.5 FL LSK subsets can be segregated into seven transcriptional clusters based on single-cell RNA-Sequencing data. Heatmap represents marker genes for each transcriptional cluster. Note the enrichment of HSC-associated genes in cluster 4 which are composed of cells from both iLT-HSC and iST-HSC populations. **(B)** Intra-cluster mean average (MA) plots of iLT-HSCs versus iST-HSCs in cluster 4 **(i)** and iLT-HSCs versus iST-HSCs in cluster 7 **(ii)**. **(C)** Inter-cluster mean average (MA) plots of iLT-HSCs in cluster 4 versus cluster 7 **(i)** and iST-HSCs in cluster 4 versus cluster 7 **(ii)**. **(D) (i)** Heatmap of gene differentially expressed between iST-HSCs in cluster 4 versus cluster 7. **(ii)** Representative flow cytometry profiles of PTPRC (CD45) expression by E14.5 FL iLT-HSCs and iST-HSCs. *n* = 14 independent embryos. Inset values represent mean ± SEM. **(E – F)** Hematopoietic reconstitution after transplantation of 100 iLT-HSC CD45^+^ cells, 100 iST-HSC CD45^+^ cells, or 100 iST-HSC CD45^low/-^ cells into lethally irradiated mice. Peripheral blood Plts, Ery, and Leuco lineages were analyzed 2 **(E)** or 16 **(F)** weeks after transplantation. Data were analyzed using one-way ANOVA. The Šídák test was used to correct for multiple comparisons. ns, not statistically significant. *, *p* < 0.05. ****, *p* < 0.00005. iLT-HSC CD45^+^, *n* = 7 independent recipient mice. iST-HSC CD45^+^, *n* = 9 independent recipient mice. iST-HSC CD45^low/-^, *n* = 8 independent recipient mice. Dashed line represents the 1% reconstitution threshold. **(G)** Secondary transplantations were performed using 1/10 doses of primary recipients that had received iST-HSC CD45^+^ cells. Primary recipients had originally received a 100-cell dose of iST-HSC CD45^+^ cells. Peripheral blood Plt, Ery, and Leuco lineages were analyzed 16 weeks after transplantation. *n* = 3 secondary recipients from 2 independent primary recipients. Inset values represent the frequency of reconstitution secondary recipients. Dashed line represents the 1% reconstitution threshold. Plt, platelets. Ery, erythrocytes. Leuco, leucocytes.

Given that LT-RU frequency in the iST-HSC population was half that observed in the iLT-HSC population, to directly compare iST:LT-RUs with iLT:LT-RU we first aimed to improve iST:LT-RU enrichment. To this end, differential gene expression analysis was performed between iST-HSCs in cluster 4 and cluster 7. Amongst the significantly upregulated genes in the cluster 4, *Ptprc* (which encodes CD45) was found to be strongly up-regulated (Figure 6Di and Table S10). Other genes included *Procr* (which encodes EPCR) and *Esam* (Figure 6Di and Table S10), both of which are known to be expressed by E14.5 iLT-HSCs and enrich for LT-RU activity (Balazs et al., 2006; Kent et al., 2009; Yokota et al., 2009). CD45 is expressed by LT-RUs once they have emerged in the E11.5 AGM region (Taoudi et al., 2008; Taoudi et al., 2005), in the E12.5 yolk sac (Taoudi et al., 2005), and in the E13.5 – E15.5 FL (Kent et al., 2009; Taoudi et al., 2005). Thus, we considered CD45 expression to be a promising marker for improved FL iST:LT-RU enrichment. Using flow cytometry we found that 85% ± 10% of iLT-HSCs and 56% ± 12% of iST-HSCs were CD45+ (Figure 6Dii).

We next compared the ability of iLT-HSCs, CD45+ iST-HSCs, and CD45low/-iST-HSCs to provide acute-and long-term reconstitution. To this end, 100 cells of each immunophenotype were transplanted into lethally irradiated recipients. In keeping with the enhanced reconstitution observed with IGF2-signalling and *Satb1* expression (Doi et al., 2018; Thomas et al., 2016; Zhang and Lodish, 2004), after two weeks CD45 low/-iST-HSCs cells failed to contribute to reconstitution but CD45+ iST-HSCs cells provided robust reconstitution that was significantly greater than that of iLT-HSCs (Figure 6E). By 16 weeks, CD45 low/-iST-HSCs did not provide any reconstitution, but iLT-HSCs and CD45+ iST-HSCs provided equivalent hematopoietic contribution (Figure 6F). Secondary transplantation demonstrated the long-term self-renewal capacity of CD45+ iST-HSC LT-RU reconstitution (Figure 6G). As all LT-RUs were contained within the 56% of iST-HSCs that expressed CD45 (Figure 6Dii), the LT-RU frequency in the CD45+ iST-HSC population in ∼ 1/19 cells, a frequency equivalent to LT-RUs in the iLT-HSC population (Figure 5D).

### The iST:LT-RU variant does not contain developmentally-restricted (lymphoid-biased) HSCs but is labelled by Flt3-cre

Developmentally-restricted HSCs (drHSCs) (Beaudin et al., 2016) are a component of the E14.5 FL LSK. Similar to LT-HSCs, drHSCs express CD150 and can self-renew in secondary transplantation assays. However, under physiological conditions drHSCs exhaust between the neonatal period and adulthood. Two features distinguish drHSCs from LT-HSCs: (1) from eight weeks post-transplantation, drHSC-derived reconstitution is lymphoid-biased (Beaudin et al., 2016). This bias is transferred upon secondary transplantation (Beaudin et al., 2016); and, (2) drHSCs emerge via a *Flt3*-expressing ancestry. Use of a *Flt3*-cre expressing mouse line elegantly distinguishes LT-HSCs from drHSCs in E14.5 FL LSK CD150+ compartment (Beaudin et al., 2016). Our data showed that iST:LT-RUs were CD150- and provided lineage-balanced long-term reconstitution, thus were not a component of the drHSC axis.

A recent study demonstrated cells that non LT-HSCs that derive via a *Flt3*-expressing ancestry contribute to native hematopoiesis in adulthood (Patel et al., 2022); these cells were termed long-lived embryonic MPPs. To investigate if iST:LT-RUs were a component of this long-lived embryo-derived axis of native adult hematopoiesis, we inter-crossed *Flt3*-cre mice with floxed-STOP *Rosa26*-EYFP mice. In keeping with the findings of Patel *et al* (Patel et al., 2022), we found that CD45+ iST-HSCs were significantly more effectively labelled than iLT-HSCs (Figure 7A). To compare reconstitution potential, 100 YFP+ or YFP-iST-HSC[CD45+] cells were transplanted into lethally irradiated mice. After 4 weeks YFP+ and YFP-cells contributed to monocyte and B-cell reconstitution (Figure 7B); after 16 weeks robust monocyte, B-cell, and T-cell reconstitution was observed (Figure 7C). Thus, iST:LT-RUs of *Flt3*-expressing ancestry were capable of long-term lineage-balanced hematopoietic reconstitution.

**Figure 7.**
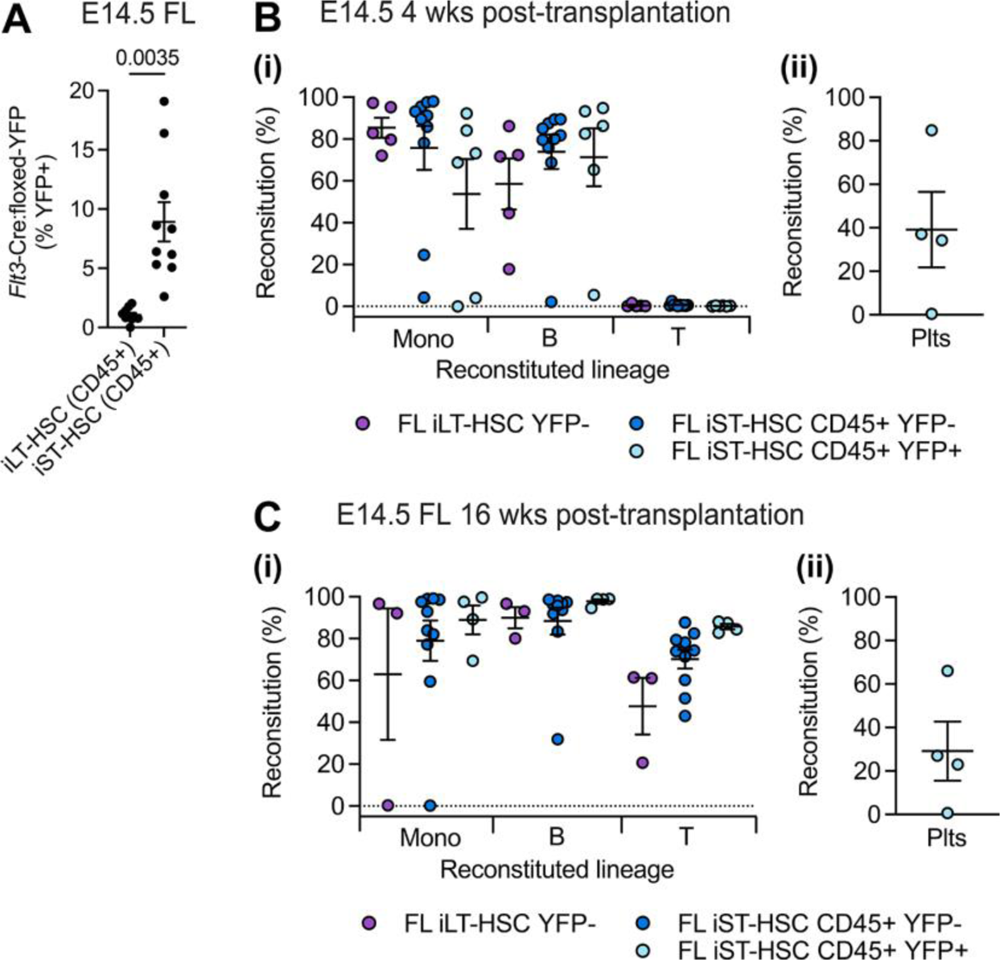
E14.5 iST-HSCs contain LT-RUs of *Flt3*-cre ancestry that have lineage balanced in vivo outputs. **(A)** Quantification of *Flt3*-cre driven recombination of the floxed-YFP locus in E14.5 FL iLT-HSC and iST-HSC CD45+ populations. *n* = 10 independent mice. Data were analyzed using the unpaired two-tailed student T-test. Exact p value is shown. **(B** and **C)** E14.5 FL HSCs were fractioned according to *Flt3*-cre:floxed-YFP labelling. 100 cells YFP+ or YFP-cells from population were injected into lethally irradiated recipients and analyzed after 4 **(B)** and 16 **(C)** weeks. Due to the paucity of iLT-HSC YFP+ cells, this fraction was not transplanted. **(i)** Contribution to **p**eripheral blood monocytes (Mono), B-cells (B), and T-cells (T). **(ii)** Reconstitution of peripheral blood platelet (Plts). Data were analyzed using one-way ANOVA. The Šídák test was used to correct for multiple comparisons. Only comparisons where *p* < 0.05 are displayed. Exact p values are shown. iST-HSC CD45+ YFP-, *n* = 10 independent recipient mice. iST-HSC CD45+ YFP+, *n* = 6 independent recipient mice.

## Discussion

Transcriptional analysis of iMPP2 – 4 from the E14.5 FL revealed that these populations were transcriptionally diverse along axes that were largely predictable based on insight from their adult counterparts (Pietras et al., 2015): iMPP2 was distinguished by the expression of megakaryocytic and erythroid genes, iMPP4 was distinguished by the expression of lymphopoiesis genes, and iMPP3 expressed a balance of megakaryocytic, erythroid, and lymphoid genes. Functional analysis (via population-level transplantation) demonstrated that iMPP2 had a platelet (megakaryocyte), erythrocyte, and monocyte production bias; iMPP4 exhibited a lymphoid bias (relative to iMPP2) and, iMPP3 was statistically similar to iMPP4. Clonal investigation revealed that the FL iMPPs were a heterogeneous mix cells that underwent single-or dual-lineage outcomes, while multi-lineage outcomes were largely reserved for the iLT-and iST-HSC subsets. The subdued clonal capacity of FL iMPPs is consistent with observation made from ABM iMPPs following transplantation (Naik et al., 2013) and during native hematopoiesis (Rodriguez-Fraticelli et al., 2018).

The most distinguishing feature of FL iMPPs was the capacity of iMPP3 to produce B-cells, this outcome largely derived from clones that were capable of monocyte and B-cell production, whereas iMPP4 achieved B-cell production predominantly via uni-outcome clones. E14.5 FL LT-HSCs and ST-HSCs were strikingly distinct from the iMPPs. This was most evident in the lentiviral barcoding experiments where complete multi-lineage outcomes were only repeatedly detected in the HSC subsets.

We made the surprising discovery that in acute-term (two weeks), short-term (four weeks), and long-term (> 16 weeks) transplantation assays the E14.5 iST-HSC population was functionally very similar to E14.5 iLT-HSCs, including clonal output. E14.5 long-term repopulating units (LT-RUs) were not only distributed between the iLT-HSC and iST-HSC immunophenotypes, but most LT-RUs were concealed within the CD45-expressing fraction of the iST-HSC population. ABM iST-HSCs have the capacity for short-term multi-lineage reconstitution but were generally less able to sustain this in the long-term, and ABM iST-HSCs generally could not reconstitute secondary recipients (thus did not self-renew) (Morita et al., 2010; Oguro et al., 2013; Pietras et al., 2015). In contrast to this, we found that LT-RUs within the E14.5 ST-HSC population (iST:LT-RUs) were functionally similar to LT-RUs within the E14.5 LT-HSC population (iLT:LT-RUs), this included: durable multilineage reconstitution for > 40 weeks post-transplantation and following secondary transplantation; and clonal multi-lineage production which enabled the production of platelet, erythroid, monocyte, and B-cell lineages following transplantation.

The E14.5 FL iLT-HSC population contains a sub-class of CD150+ LT-RUs known as developmentally-restricted HSCs (drHSCs) which are lymphoid-biased HSCs that contribute to the first HSC-derived lineages of the fetus (Beaudin et al., 2016). Using the *Flt3*-cre:YFP mouse line (which labels drHSCs (Beaudin et al., 2016)) we found that the FL iST-HSCs were significantly more labelled than iLT-HSCs, and that *bona fide* LT-RUs within the iST-HSC had derived via a *Flt3*-expressing ancestry. This suggests that multiple LT-RU forming pathways exist to establish the E14.5 fetal stem cell pool: one that derives via a *Flt3*-expressing ancestry that yielded YFP+ LT-RUs in the iST-HSC population and one that yielded LT-RUs in iLT-HSC that were predominantly YFP-. Whether these pathways originate from different anatomical sites, or from different precursors within the same stem-cell forming organ will be a subject of future investigation.

Taken together with previous studies, our findings indicate that three LT-RU populations must co-exist in the E14.5 fetal liver: conventional CD150+ iLT:LT-RUs (Kim et al., 2006), CD150+ drHSCs (Beaudin et al., 2016), and CD150-iST:LT-RUs. Although other studies have previously shown that LT-RUs were present in the CD150-fraction of the E14.5 FL (Kent et al., 2009; Kim et al., 2006; Papathanasiou et al., 2009) they did not quantify the LT-RU biomass distribution between the CD150+ and CD150-fractions of LSK FLT3-CD48-cells. Thus, the significance of CD150-LT-RU went unappreciated. We found that approximately 70 % of E14.5 FL LT-RUs are within the ST-HSC immunophenotype (LSK_FLT3-CD150-CD48-, Figure 5E). Based on our findings (analysis of our scRNA-Seq data and functional tests), the use of LT-RU-associated markers such as EPCR (Balazs et al., 2006; Kent et al., 2009), ESAM (Yokota et al., 2009), or CD45 ((Kent et al., 2009; Taoudi et al., 2008; Taoudi et al., 2005) and Figure 6E – G) in LSK_FLT3-CD48-cells would provide a single unifying LT-RU enriching immunophenotype. However, this approach would censor important information such as differences in the developmental pathways of iLT-HSCs and iST-HSCs as indicating by *Flt3*-expressing ancestry ((Patel et al., 2022) and Figure 7).

When LT-RUs first emerge in the E11.5 AGM region they do not express CD150 (McKinney-Freeman et al., 2009), nor do LT-RUs express CD150 in the E12.5 placenta (McKinney-Freeman et al., 2009). Although E14.5 LT-RUs can be purified at high frequency according to CD150 expression (Kim et al., 2006), our data demonstrates that the E14.5 FL LT-RUs biomass is a mix of CD150- and CD150+ LT-RUs. We were not able to test whether during continued physiological development E14.5 FL iST:LT-RUs would initiate CD150 expression and so become iLT:LT-RUs. Although it could be presumed that early LT-RUs (e.g, those produced in the AGM region) all upregulate CD150 expression during fetal development, this is currently untested. Of note, findings from a previous study suggest that FL iLT-HSCs and iST-HSCs are ontologically distinct lineages that largely maintain segregation and persist into adulthood: using a *Flt3*-creERT2 mouse line, Patel *et al* (Patel et al., 2022) demonstrated that after inducing labelling at E10.5 or at E14.5, iST-HSCs and iMPPs in the E15.5 FL could be labelled without labelling iLT-HSCs. The authors noted that *Flt3*-creERT2 labelled lineages from the fetus persisted into adulthood and contributed to ongoing hematopoiesis (Patel et al., 2022). Although it was concluded that embryonic MPPs drove the persistence of life-long *Flt3*-cre derived hematopoiesis, given our findings it is possible that LT-RUs present in the iST-HSC population could persist into adulthood and contribute to native hematopoiesis.

## Acknowledgments

This work was supported by: the Australian Research Council (ARC) Stem Cells Australia program; NH&MRC project (1128993, 1129012, 2011770), program (1113577), and Investigator (1173342) grants; Independent Research Institutes Infrastructure Support Scheme grant 361646 from the NH&MRC, the Australian Cancer Research Fund, and Victorian State Government Operational Infrastructure Support. S.T was supported by a fellowship from the Lorenzo and Pamela Galli Charitable Trust and a Royal Society Wolfson Fellowship (233002).

## Author contributions

O.J.S designed experiments, performed experiments, analyzed data, and wrote the manuscript first draft. C.B designed experiments, performed experiments, and analyzed data. A.G analyzed scRNA-Seq data. K.A.F, M.A.D, and S.H.N generated essential reagents and designed experiments. T.S.W analyzed lentiviral barcoding data. S.T, A.F, A.F.T, and W.S.A provided essential intellectual input. S.T. conceived the study, designed experiments, and analyzed data. All authors contributed to writing the manuscript.

## Materials and correspondence

Correspondence should be addressed to Samir Taoudi.

## Supplementary information

### Supplementary Figures

**Figure S1.**
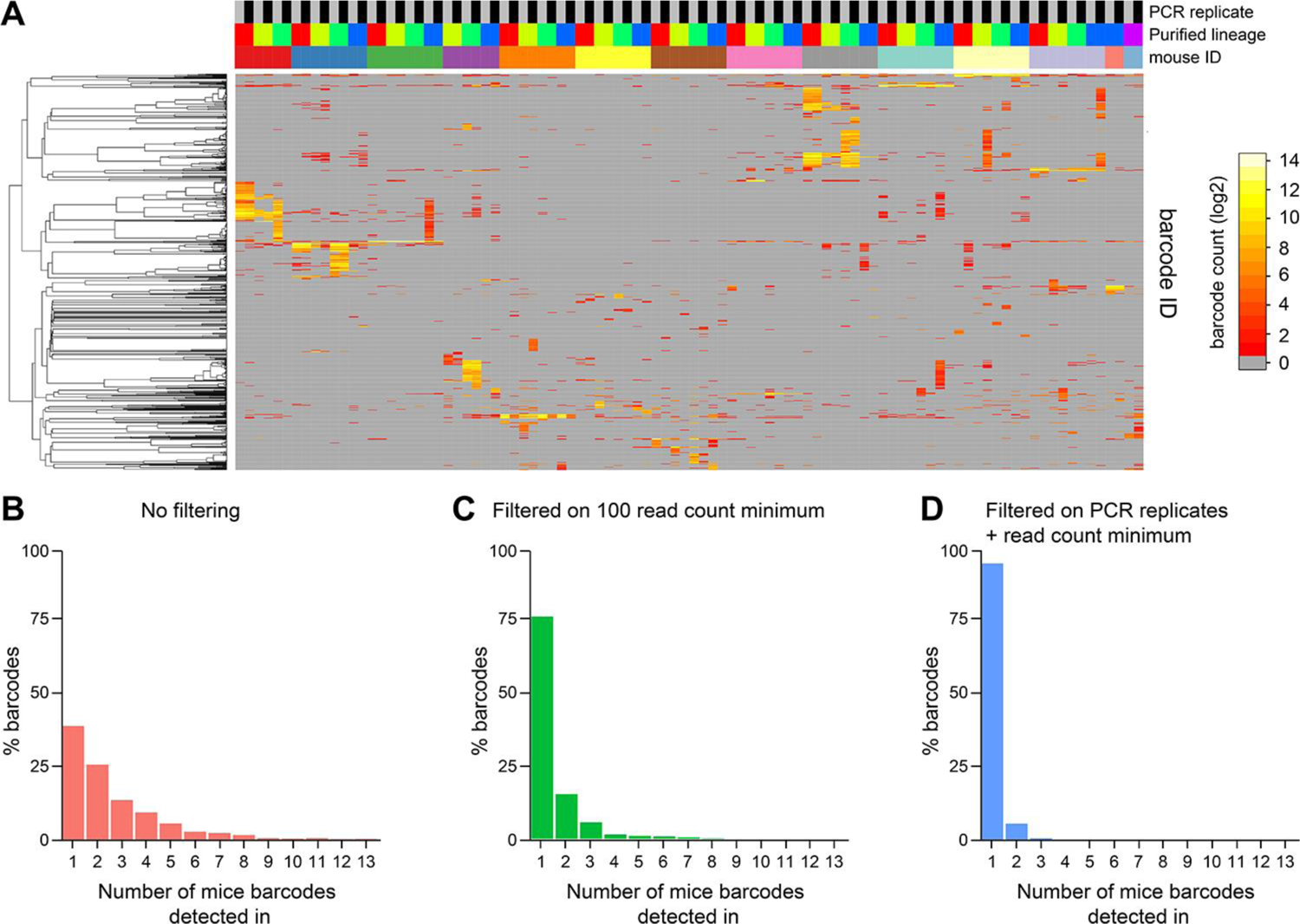
Filtering lentiviral barcoding sequences. **(A)** Heatmap of all barcode sequences detected before filtering in the experimental cohort of 14 animals. Mouse ID are color-coded. Purified lineage: Red: B cells, Yellow: Erythrocytes, Green: Monocytes, Blue: Neutrophils, Purple: blank. 2 PCR replicates (grey/black) were performed for each sample. **(B – D)** Barcode sharing between all cohorts of experimental mice with no data filtering **(B)**, after removal of low read counts (“min_read”) **(C)**, and after PCR replicate filtering (“PCR rep”) and “min_read” filtering **(D)**. Data analyzed in this study were based on “PCR rep” + “min_read” filtering. This shows that >95% bona fide barcodes are detected in one mouse only, suggesting that repeated usage of barcodes is unlikely.

**Figure S2.**
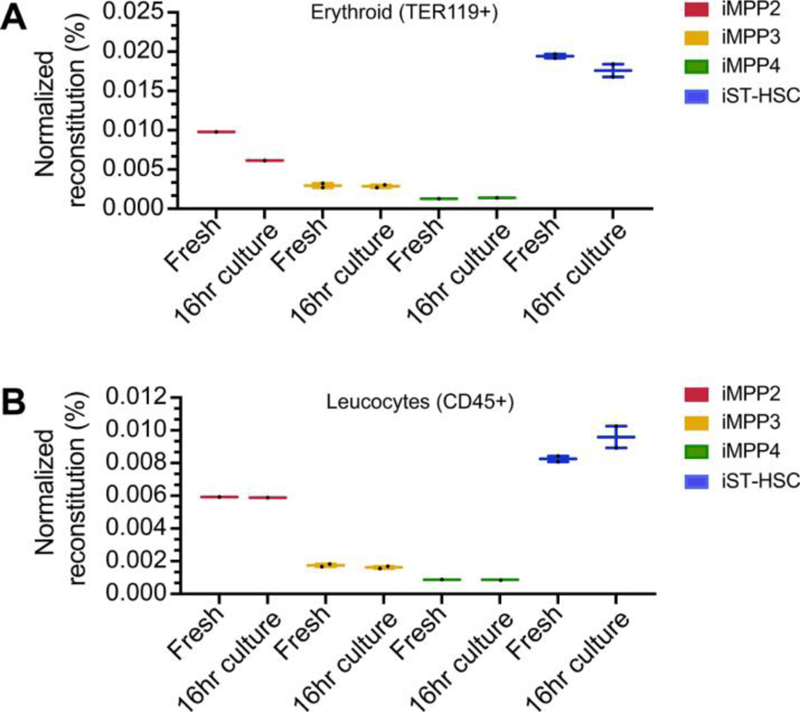
16 hrs of *ex vivo* culture did not affect LSK subset function. Normalized reconstitution per cell two weeks after transplantation of freshly isolated E14.5 FL LSK subsets or E14.5 FL LSK subsets following 16 hrs of *in vitro* culture. **(A)** Reconstitution in the erythroid lineage. **(B)** Reconstitution in leucocytes. iMPP2 *n* = 1 recipient. iMPP3, *n* = 2 independent recipients. iMPP4, *n* = 1 recipient. iST-HSC, *n* = 2 independent recipients.

**Figure S3.**
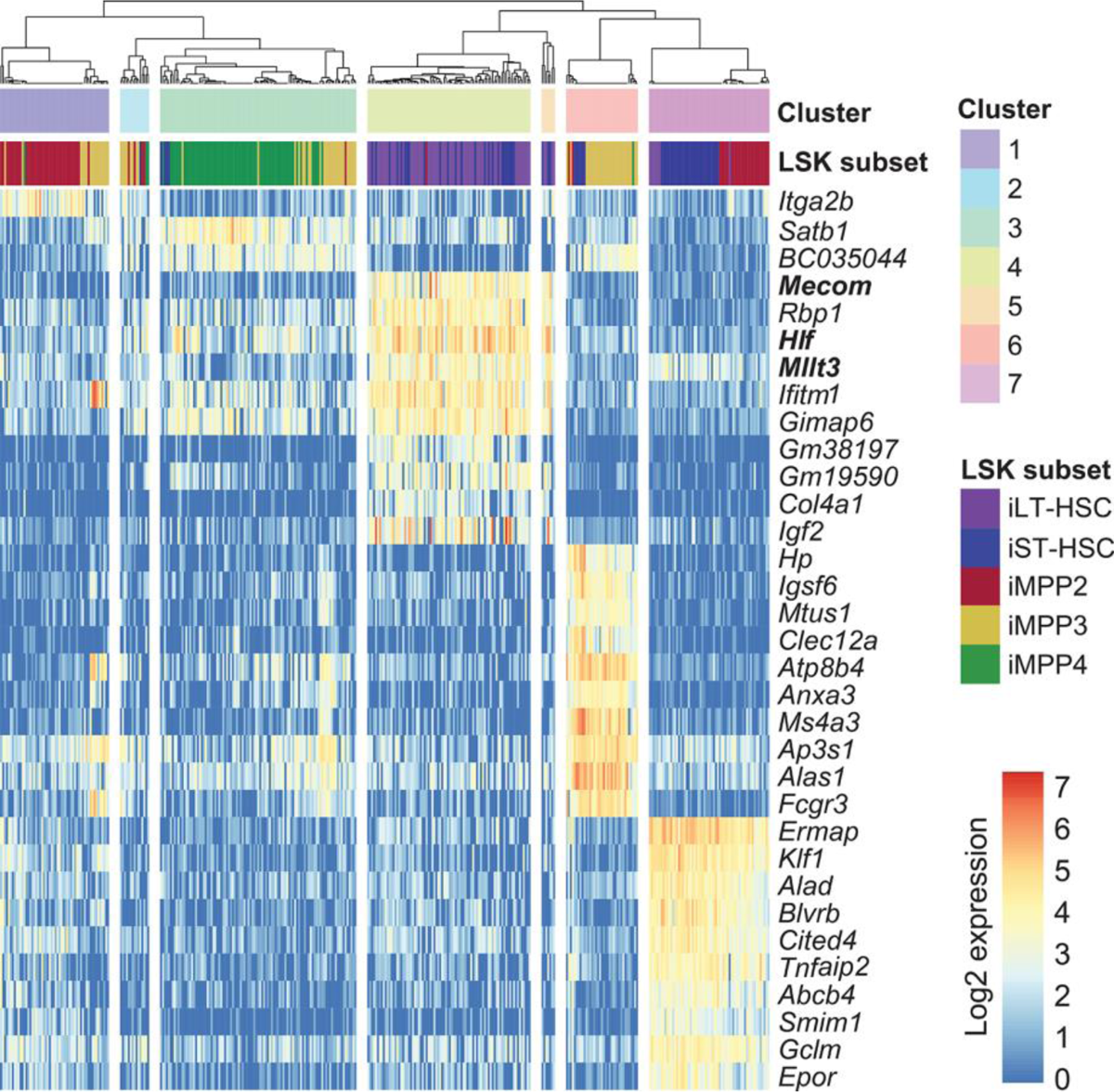
Signature genes of E14.5 FL LSK transcriptional clusters. Heatmap of E14.5 FL LSK transcriptional cluster signature genes (linked to Figure 6A).

